# Transferring structural knowledge across cognitive maps in humans and models

**DOI:** 10.1101/860478

**Authors:** Shirley Mark, Rani Moran, Thomas Parr, Steve Kennerley, Tim Behrens

## Abstract

Relations between task elements often follow hidden underlying structural forms such as periodicities or hierarchies, whose inferences fosters performance. However, transferring structural knowledge to novel environments requires flexible representations that are generalizable over particularities of the current environment, such as its stimuli and size. We suggest that humans represent structural forms as abstract basis sets and that in novel tasks, the structural form is inferred and the relevant basis set is transferred. Using a computational model, we show that such representation allows inference of the underlying structural form, important task states, effective behavioural policies and the existence of unobserved state-trajectories. In two experiments, participants learned three abstract graphs during two successive days. We tested how structural knowledge acquired on Day-1 affected Day-2 performance. In line with our model, participants who had a correct structural prior were able to infer the existence of unobserved state-trajectories and appropriate behavioural policies.

## Introduction

Decisions in a new environment require the understanding of what are the relevant components in this environment and how they are related to each other. In the cognitive literature, the representation that holds such information is termed a ‘cognitive map’ ^1^. Equipped with a ‘cognitive map’, an animal can predict the consequence of events and actions to inform its decisions. In novel environments, while learning a new cognitive map, it should be beneficial to exploit relevant information that was acquired in the past. What information is relevant to transfer and how it is represented is still an open question.

One possibility is that, whilst sensory information may differ radically between different situations, the brain may take advantage of previously learnt *structural* knowledge. Relationships between elements in different environments often follow stereotypical patterns ^2–4^. Social networks, for example, are organized in communities ^5^. The day-night cycle, the cycle over the seasons and the appearance of the moon in the sky all follow a periodic pattern. Hierarchies are also abundant, for example, a family, management chain in a workplace or concept organization ^6^. Representing such structures confers theoretical advantages in learning when encountering a new environment. Inferring the relevant structure enables the use of policies that are beneficial in environments with a particular underlying structure. Further, relationships that have never been observed can be inferred because the structure of the problem is familiar ^7–10^.

One structure which is pervasive in life and which *we know* facilitates such inferences is the 2-dimensional topology of space. This structure can be used, for example, to infer the correct trajectory to a goal even when the intermediate locations have never been experienced (as with ^11,12^). If such structural knowledge can be transferred from one set of sensory events to another, it should be represented in a way that is disentangled from the sensory stimuli and the particularities of the current task. We can think of representing all tasks as graphs, each node on the graph is a particular sensory stimulus that is currently experienced, for example, observing the shape of the moon. Then, an edge between two sensory stimuli implies a transition between sensory states; a round moon will be followed by an elliptic moon. These graphs can have different structural forms ^13,14^. The lunar graph, the seasonal graph and the day-night graph will all be circular; a work place graph will be hierarchical; the social network graph will have a community structure; and the spatial environment will have a transition structure that respects the translational and rotational invariances of 2D space. Can humans extract such abstract information and use it to facilitate new inferences? If so, how can this knowledge be represented efficiently by the brain?

Here we show that humans extract structural regularities in graph-learning tasks. When observing a new sensory environment with a familiar structural form, they infer the existence of paths they have never seen that conform to the structural form and make novel choices that are likely beneficial. Furthermore, participants change their strategy for learning a new graph to match previously experienced graph statistics. In order to understand these effects, we suggest a computational mechanism for representing this structural knowledge. Such representation should allow inference of the currently relevant structural form and the transfer of relevant knowledge to new sensory environments. Structural abstraction is inherent to common computational frameworks such as Hidden Markov Models. However, for flexible generalisation to new environments, the representation should highlight key statistical properties of the graph structure but suppress environment-specific particularities. We show that this can be achieved by representing structural knowledge in the form of basis sets (a set of vectors that can be used for function approximation), as has been proposed in reinforcement learning ^8,15^. This complements generative modelling approaches that attempt to infer low-dimensional latent states that explain high dimensional observations ^16^, and complies with the Bayesian Occam’s razor in finding the simplest explanation for these ^17^.

## Results

We created a task in which simulated agents and humans learn abstract graphs (Figure 1). The graphs belong to two different structural forms. Each structural form is controlled by a different connectivity rule (Figures 1A). We focused on two structural forms, a graph with a transition matrix that obeys translational and rotational invariant symmetry (Hexagonal graph) and graphs that have underlying community structure (Figures 1A). Each node on the graph corresponds to a sensory stimulus (a picture - which is a state of the task). Each edge implies that these sensory stimuli can appear one after the other (direct bidirectional transitions between the states are allowed ^18,19^). Using this task, we asked whether our agents and participants can infer the structural form of the underlying graph and how the structural knowledge can be exploited to better accomplish the task. Participants performed the task during two successive days. We asked whether participants can infer the underlying structural form during the first day and transfer and exploit this knowledge during the second day.

**Figure 1:**
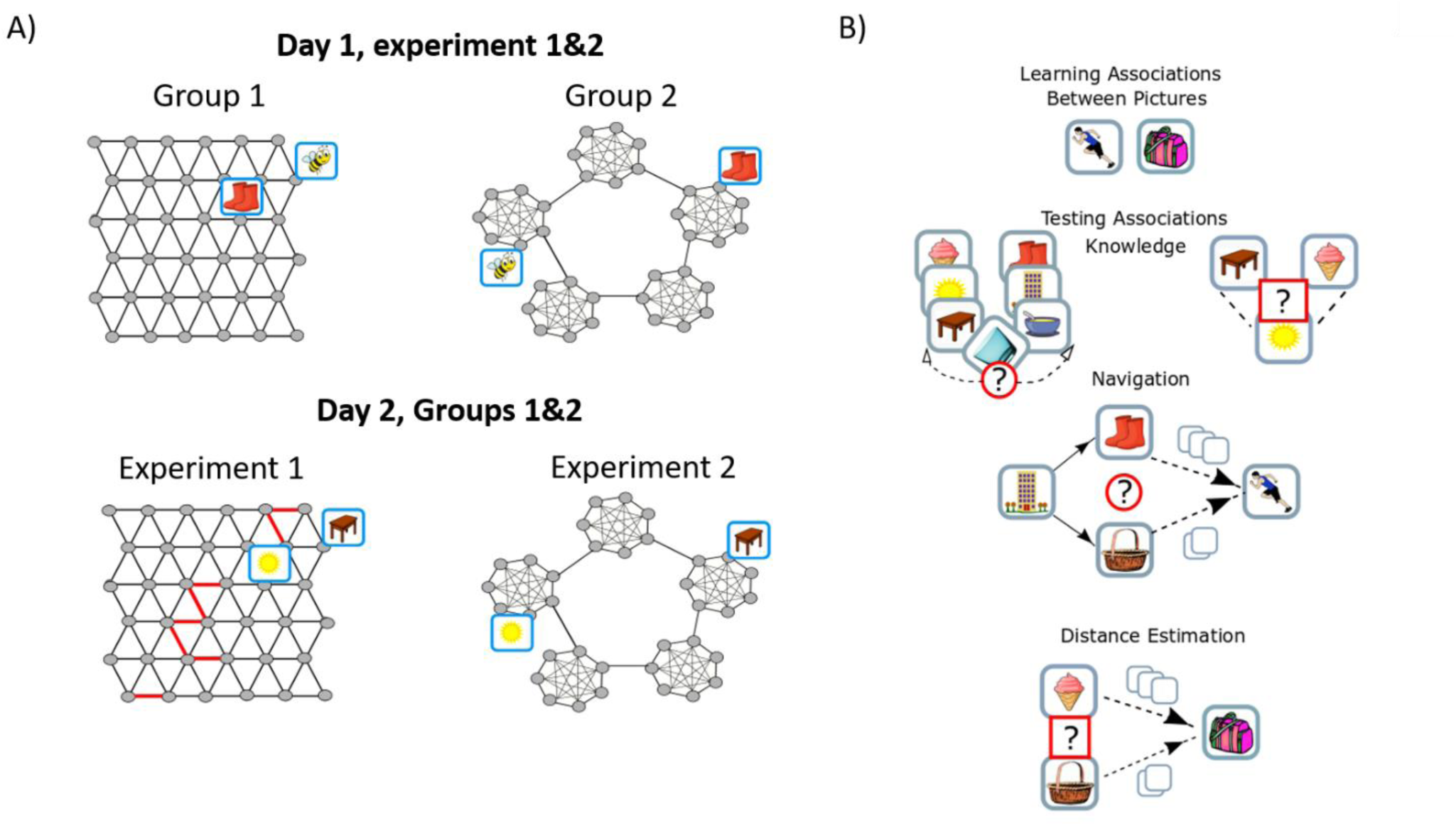
Transfer of structural knowledge: Graph structures and experimental design. A) Experimental design. Agent and participants learned graphs with underlying Hexagonal (left) or Community (right) structure. Each grey dot is a node on a graph and corresponds to a picture that was viewed by the participant (for example, a picture of a bee). The lines are edges between nodes. Pictures of nodes that are connected by an edge can appear one after the other. The degree of all nodes in both graphs is six (a connecting node connects to one fewer nodes within a community to keep the degree six). Participants learned the graph during two successive days. In both experiments, participants were segregated into two groups. Participants of one group learned graphs with the same underlying structure during both days while the other groups learned graphs with different underlying structures during the different days. B) One block of the task. Participants never observed the underlying graph structure but had to learn (or infer) it by performing a task. On each block, participants learned the associations between pairs of pictures, each pair of pictures are emitted by neighboured states on the graph. Following the learning phase, they had to answer different types of questions: 1) report which of two pictures sequences could be extended with a target picture. 2) indicate whether the picture in the middle (sun) can appear between the two other pictures in a sequence (left and right, respectively, under “testing Associations Knowledge”) 3) they navigated on the graph: starting from a certain picture, for example, the building, they had to choose (or skip) between two pictures that are connected to the current picture (the building) on the graph, for example, basket and boots (empty squares above the arrows indicate minimum steps), the chosen picture then replaces the ‘starting picture’. They had to repeat this step until they get the target picture (for example the running man). 4) which picture is closer to the target picture (the bag), the ice-cream or the basket? (Distance Estimation).

The agents and participants learned the graphs by observing pairs of stimuli that are connected by an edge. Each block of the task starts with a learning phase, following the learning phase, we examined agents’ and participants’ knowledge of the graph. To examine participants’ knowledge of the graph, we performed four separate tests within each block of the task (Figure 1b): 1) We ask participants to report which of two picture sequences could be extended with a target picture. (2) Participants reported whether a target picture could appear between two other pictures in a sequence 3) Participants navigated on the graph; starting from a source picture, participants repeatedly chose between two of the picture’s neighbours until reaching the target, with the aim to do so in the smallest number of steps. 4) Participants were asked to report which of two pictures is closer to a target picture (without feedback).

We tested transfer of structural knowledge by conducting two different experiments. On each experiment, we tested the effect of transfer of a specific structural form; on the first experiment, we have tested transfer of Hexagonal grid structure, while in the second we have tested the transfer of community structure knowledge. In each experiment, participants learned two graphs with a particular structural form during the first day. The effect of prior structural knowledge was then tested on the following day by examining participants learning of a third graph (Figure 1). To test for transfer of structural knowledge, in each experiment, we divided participants into two different groups. One group was exposed to graphs with the same structural form (but different images) on both days. The second group was exposed to graphs with different structural forms (and images) on each day (Figure 1A). This design allows us to control for all effects that are independent of the structure of the graphs; as the task is independent of the structural form, and its identity is not explicitly observed by the participants.

### Inferring and transferring graph structure

In order to better understand the problem, we considered a model of this task. The task of the participants and agent is to learn the graph. One solution to such a problem is to represent all the associations between the stimuli ^18,20^ (Figure 2, left). This results in conjunctive representation of the stimuli and their relationships. Although this type of representation can be useful, it does not allow generalization, as it is tied to the particular stimuli.

**Figure 2:**
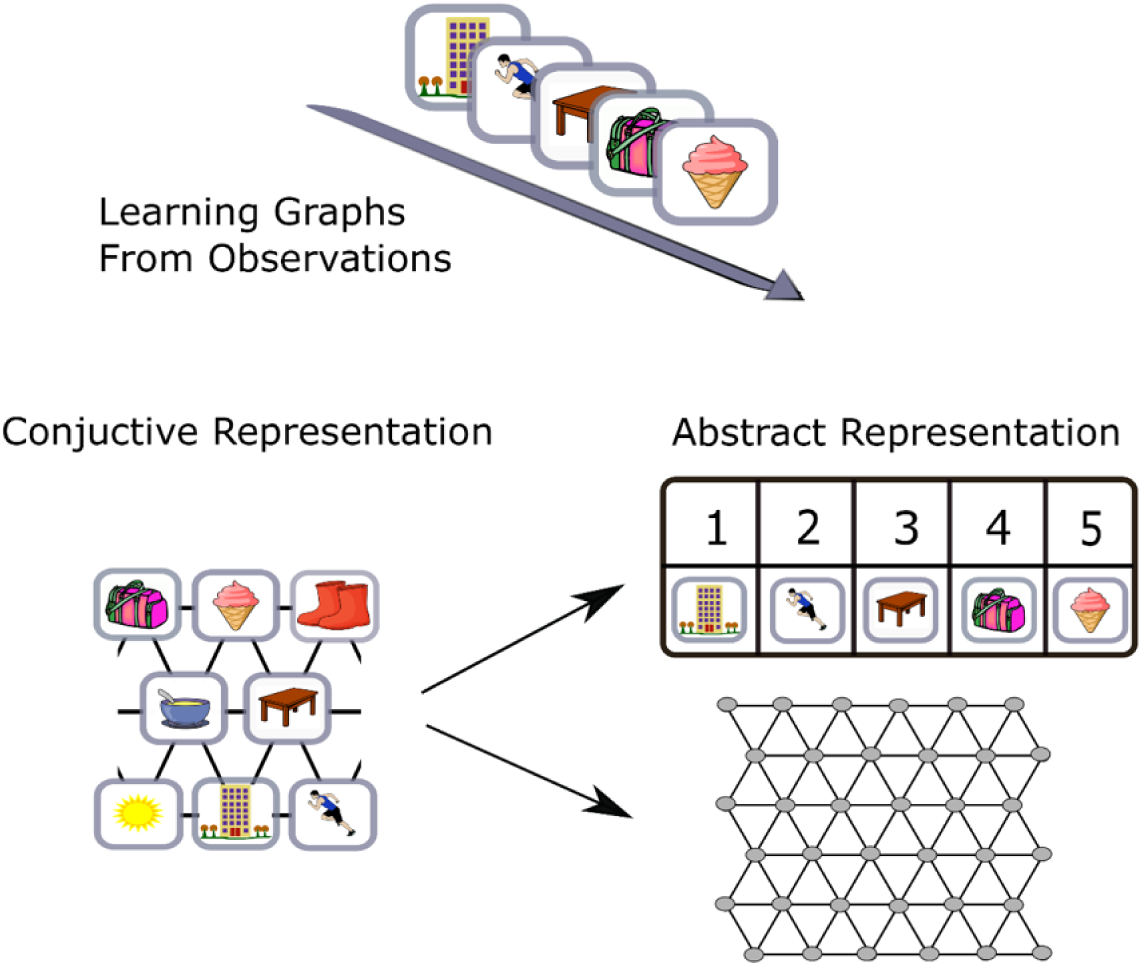
Explicit and abstract representation of transition structure. Learning underlying graph structure from observation of pictures. Graph can be represented by learning the associations between the stimuli. Such learning results in conjunctive representation that does not allow generalization and knowledge transfer. Representing the graph using two separate matrices, the transition and emission matrices allows generalization over graph structure.

Instead, we considered a Hidden Markov Model (HMM). The Markov assumption is that each latent state depends directly only on the state at the previous time-step. In other words, the past is independent of the future conditioned on the present. This dependency is captured by the transition matrix, *A*. Each entry, *Aij*, in this matrix represents the probability to move from state *i* to state *j*. A second matrix, the emission matrix *Bik*, represents the probability that state *i* will emit observation (particular stimulus) *k*. Together, both these matrices describe the probability of a sequence of observations. The HMM framework is promising because it maintains separate representations of transitions and emissions and therefore can easily generalise transition structures to new sensory stimuli (Figure 2). However, we consider two extensions to vanilla HMMs.

First, because transition matrices (*A*) of the same structural form may not be identical (e.g. different number of nodes), we need a flexible representation of transition structure. We approximate the transition structure using basis sets for structural knowledge (Figure 3B,C &D, see below and methods for further details). Second, we propose a method for inferring, amongst candidate basis sets, the one that best fits the current task. We assume that each common structural form is represented by a basis set and this basis set is known to the agent. The agent needs to infer the underlying structural form and exploit the knowledge of the basis set to estimate the transition matrix of the current task. Once the structural form is inferred, the size of the graph should also be inferred. The agent should then adjust the basis set according to the inferred graph size and estimate the transition structure of the current task (Figure 3D, see methods for details).

**Figure 3:**
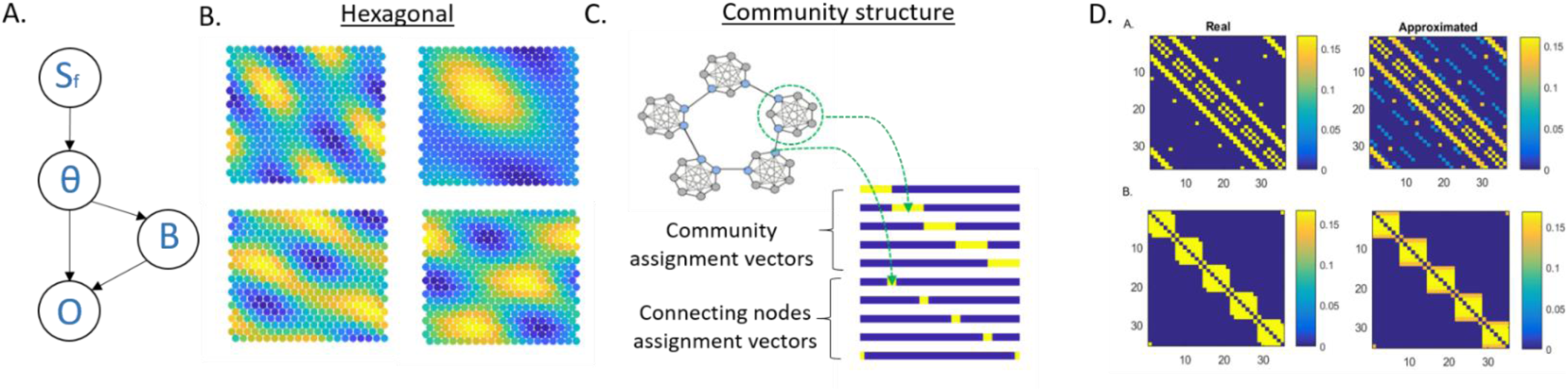
Inferring graph structure rather than learning it using a basis sets representation for structural knowledge. A) We present a generative model for graphs. Each graph belongs to a structural form (Sf). Given a structural form, graph size (θ) is sampled from prior distribution (p(θ|St)) and the transition matrix is approximated. Given a transition matrix (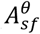, that determined by the form and the dimensions) an emission matrix (B) is sampled. From these two matrices the sequence of observations (O) can be generated. B) Basis sets for Hexagonal grid, few examples. C) Basis sets for a community structure. Basis sets can allow direct inference of important graph states without the need of further computation. In a graph with underlying community structure, the connecting nodes (blue circles) are important; knowing them allows fast transitions between communities. A basis set that contains explicit connecting nodes assignment vectors allows the direct inference of their identity by learning the emission matrix. D) The transition matrices can be approximated using Basis sets for structural knowledge. Upper panels: correct and approximated transition structure for Hexgonal grids with 36 nodes. Lower panels: real and approximated transition matrices for a graph with underlying community structure (35 nodes).

The problem at hand can be formulated as a hierarchical generative model of graphs ^21^. Each structural form, using the basis set representation, can generate a particular transition structure according to a vector of parameters (θ) that defines the particularities of the current graph, such as size. Together with the emission matrix (B), the observation (O) can be generated (Figure 3A). The job of the agent is to infer the current structural form (*S*_*f*_), graph size and the emission matrix. The agent first uses approximate Bayes (see methods) to infer the structural form and graph size:

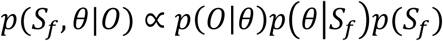

Where:

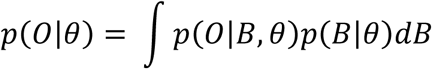

While doing so, the agent calculates the maximum likelihood estimates of the emission matrices for each considered graph (see methods). The emission matrix that corresponds to the inferred structural form and graph size is then chosen to represent the current task. Using this method, the agent was able to infer the correct structural form (Figure S1&S2) and graph size (Figure 4A, Figure 5A). Following the estimation of transition and emission matrices, the agent can estimate the distances (number of links) between observations (see methods). Indeed, when asking the agent to report which of two pictures is closer to the target picture, similarly to participants, the agent was able to perform it correctly (Figure 4A).

**Figure 4:**
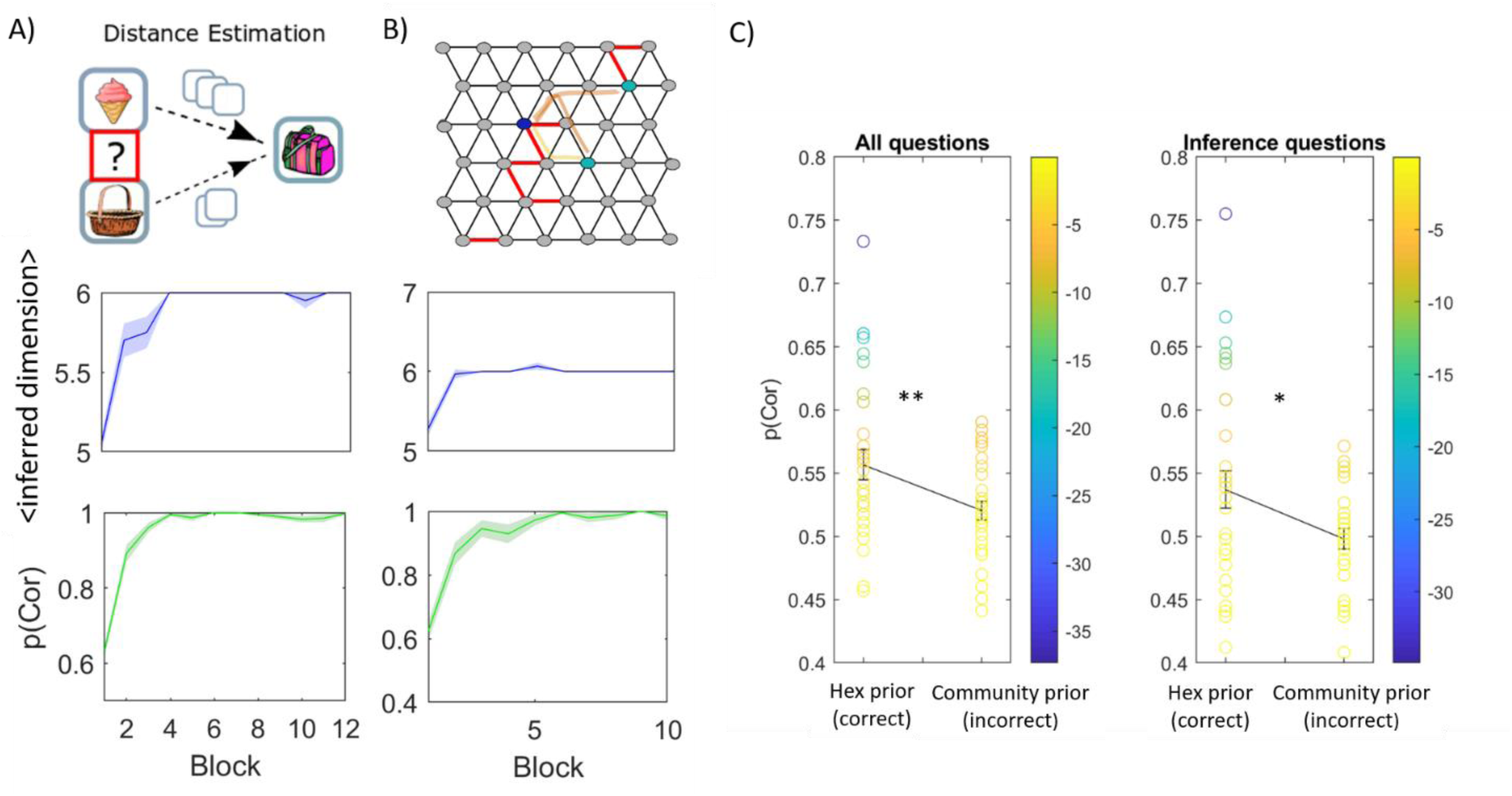
Transfer of structural knowledge allows inference of unobserved links (Hexagonal graph) A) Learning from random walk on hexagonal graph, our agent was able to infer the correct structural form and size (upper panel, Y-axis is the inferred number of nodes in one grid dimension, averaged over 15 simulations). The agent determines correctly which of two pictures is closer to a target picture using the approximated transition matrix (lower panel, y- axis is the average fraction of correct responses, out of 60 questions in each block, over 15 simulations). B) When learning from pairs that were sampled randomly (not in succession) while some of the links (pairs) were never observed (upper panel: red are links that are missing), our agent was able to infer the graph size correctly (upper panel). Further, the agent was able to infer the existence of links that were never observed and determine correctly which of two pictures is closer to a target picture, according to the complete graph. The agent could do so even though the number of observed links between the two pictures and the target was identical (p(cor) corresponds to the average fraction of correct answers out of 40 questions in each block). For example. the two nodes that are marked with light blue have the same number of observed links to the target node (marked with dark blue circle), while the number of links that connects these two nodes to the target is different on the complete graph. C) Participants had to indicate which of two pictures is closer to a target picture. Participants that reach the second day of our task with the correct prior expectation over the structural form performed significantly better in such task compared to participants with the wrong structural prior (left panel). (30 participants in each group). They were able to answer these questions significantly above chance even when there were links that were never observed, and they had to choose between two pictures with identical number of observed links to the target (right panel). Error bar: SEM.

**Figure 5:**
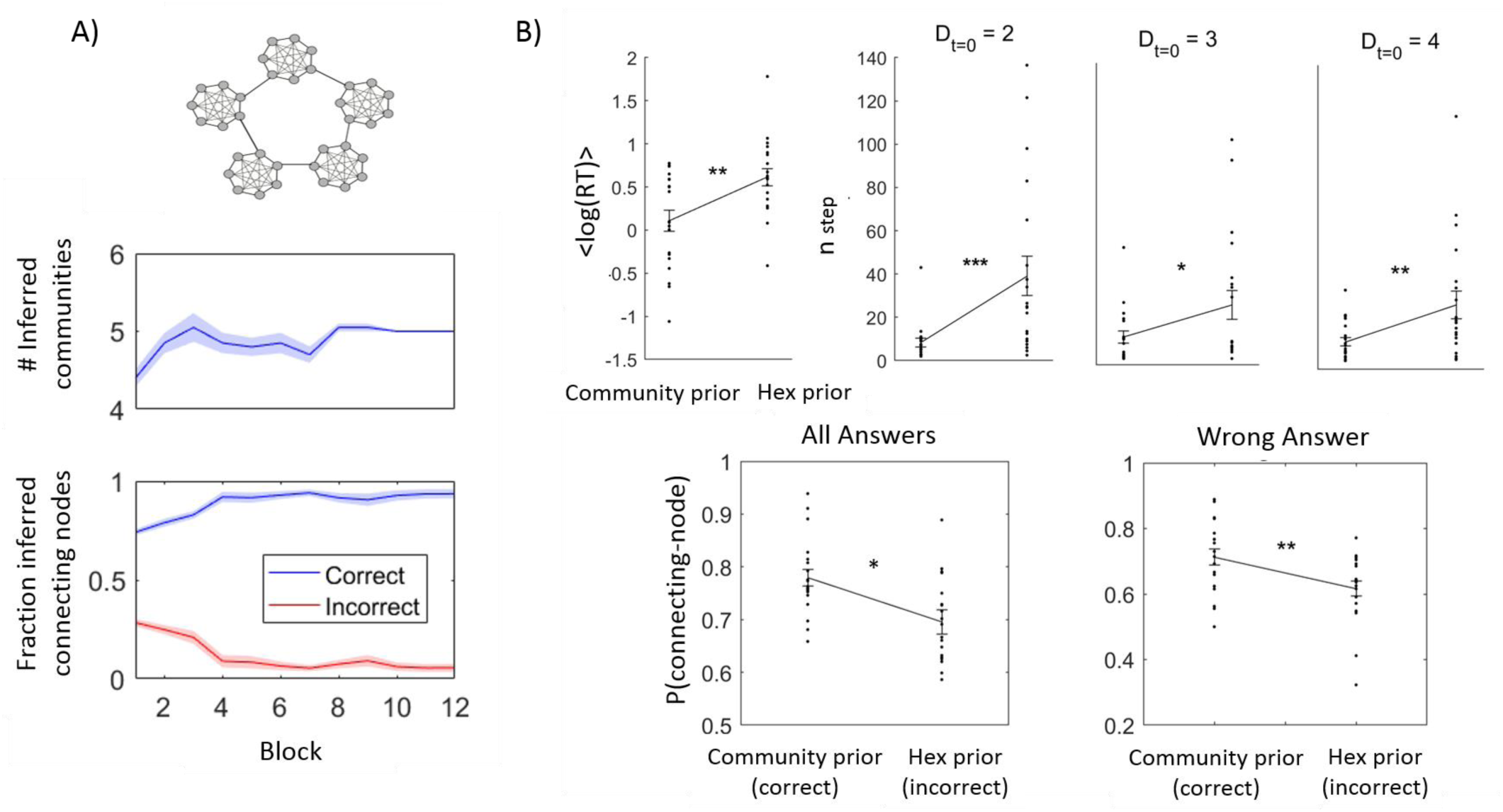
Policy transfer: Learning graphs with underlying community structure. A) Our agent was able to infer the correct number of communities (middle panel, averaged inferred number of communities over 20 simulations). It was also able to infer the identities of the connecting nodes (lower panel, inferred number of nodes divided to the number of connecting nodes according to the inferred graph size, see methods). B) Participants with correct structural prior spend less time on learning the associations between the pictures, (RT = Response Time for changing to the next pair, upper panel – left). The number of steps to the target (n_steps_) is significantly lower for participants with the correct structural prior (upper panels,D_t=0_ is the initial number of links between the current picture and the target). During navigation participants with the correct prior over the structural forms choose connecting nodes more frequently (lower panel - left), they do so even if this choice take them far away from the target (lower panel - right). (20 participants in each group)

### Basis sets definition

For basis representations of the hexagonal structure, we choose eigenvectors of a hexagonal graph transition matrix (see methods, figure 3B). These eigenvectors have previously been shown to resemble Entorhinal cortex grid cells ^22,23^. Because the community structure graph is not translationally invariant, it can be more compactly represented using bases that take two distinct forms – a set of bases for community membership and a set for connecting nodes, where ‘connecting nodes’ refers to nodes that connect two communities (Figure 3C). As we will show, this has the additional benefit of rapidly inferring connecting nodes which lead to behavioural advantages.

### Inferring unobserved trajectories

To test inference of unobserved links, we tested the model (and later the human participants) on a more difficult problem. Here, instead of a random walk, the model learns the graph by pseudo- random sampling of pairs of adjacent states. This is a harder problem than inferring the graph from random walks, as loop-closures (when the participant sees an entire trajectory returning to the same state) are far less frequent so the number of possible graphs consistent with the observations remains high for longer. However, this allowed us to perform a key manipulation. We could selectively omit key edges in the graph (red edges Figure 4b) without changing the local sequence statistics (because there were no sequences, only pseudo-random presentations of adjacent pairs). We could then ask if the agent (and later humans) could infer the existence of these omitted edges.

To test whether the model (and later humans) could infer unobserved links, we asked questions of the following form: Which of two observations is closer to the target state? In each case, the two observations were the same distance to the target, given the observed edges, but one of the observations would be closer if the model (or human) had inferred the existence of the omitted edges. The model was still able to infer the correct structural form (Figure S1) and graph size (Figure 4b). Moreover, our model was able to infer these unobserved links better than chance and answer the questions above correctly (Figure 4b, p<<0.001, see also Supplementary Figure 1). Hence, we can conclude that basis sets, as a compressed representation of transition structures, allow estimation and inference of the currently relevant transition structure and therefore enable the prediction of links that were never observed.

### People can use structural knowledge to infer unobserved trajectories

Can humans use prior knowledge of the underlying graph structure to infer the existence of transitions that were never observed? We performed graph-learning experiments where participants learned three large graphs (36 nodes with degree of 6, Figure 1a). We tested whether participants can infer (or learn) the underlying graph structure and apply this knowledge to a new graph with new stimuli. Participants were segregated into two groups. They performed the task on two successive days (Figure 1a). During the first day, one group learned two graphs with an underlying hexagonal structure while the second group learned two graphs with an underlying community structure. On that day, the graphs were learnt by observing a sequence of pictures that are taken from a random walk on the graphs.

We hypothesised that the experience during the first day shaped the prior expectations over the underlying structural forms during the second day, as participants associated the experienced graph statistics with our task. Participants who learned hexagonal graphs during the first day should expect a hexagonal graph on the second day, while participants who previously learned graphs with underlying community structure should expect to learn again a graph with a community structure. We therefore asked whether participants can infer the underlying structural form during the first day and then use it as a prior knowledge during the second day. Notably, if they do, they will be able to infer the existence of transitions they have never observed (as in the model). Therefore, as with the model above, both groups of participants learnt hexagonal graph on the second day by observing pairs of adjacent pictures. As with the model, pairs were sampled pseudo-randomly (i.e. neighbouring pairs were not sampled in succession) and many pairs were omitted. That is, many transitions were never explicitly observed by the participants (depicted in Figure 4b – red lines). We aimed to test whether participants could use structural knowledge from the first day to infer the existence of these unobserved transitions.

We used the exact same testing procedures as with the model above to examine participants’ ability to infer the existence of a link that was never observed explicitly; participants had to indicate which of two pictures is closer to a target picture; no feedback was given for this type of questions (more than 200 questions for each participant). As with the model, the two pictures were the same distance to the target, given the observed links, but one was closer if the existence of the missing link was inferred. Only participants who were able to complete ‘missing links’ using knowledge of the underlying graph structure could answer these questions correctly. Indeed, participants who had experienced the hexagonal structure on different graphs the previous day, perform significantly better than control participants who had experienced graphs with underlying community structure (Figure 4c, left: all questions, right: ‘missing links’ questions only. <Phex(cor)>= 0.54,<Pcl(cor)> = 0.5, t = 2.29, *p-value=0.013 for inference questions*, <Phex(cor)>= 0.56, <Pcl(cor)> = 0.52, t = 2.55, *p-value=0.0068 for all questions, df = 58, one-tailed ttest*). These results indicate that, similarly to our model, participants extract sophisticated structural knowledge of the problem that generalises across different sensory realisations. They were able to transfer knowledge from one day to the other and use this knowledge to guide their decisions and infer unobserved trajectories.

Notably, this effect is driven by a subset (1/3) of participants (Figure 4c). This subset performs the inference extremely reliably (individual p-values < 10^-10). We conclude that despite the group wise significant effect, only a subset of participants transferred structural knowledge of the hexagonal graph. This may be because of the difficult nature of the graph tasks, with many states, no visual or border cues to help define the graph, and no feedback. Nevertheless, participants who did infer the structure could use it to perform inference at levels far above chance. These effects cannot be driven by non-inferential approximations of graph distance (such as the successor representation ^23,24^) as all such measures consider each graph independently and are therefore invariant to the structural form of the previous day’s graph (with different stimuli).

### Using structural knowledge to set advantageous policies

Not only can structural knowledge be used to infer unobserved transitions, it can also be used to direct advantageous policies. For example, while navigating on a graph with a community structure, agents with no structural knowledge will spend large periods trapped in a single community. A simple policy of “prefer connecting nodes” overcomes this problem. When the correct underlying structure is of community structure, our model can infer it correctly (See Supplementary figure 2). Our model also infers the number of communities correctly (Figure 5a – upper panel). Using this particular basis set for transition matrix estimation allows direct identification of connecting nodes (Figure 3C). Indeed, the identity of the connecting nodes is recovered correctly during the learning of the emission matrix (Figure 5a –low panel, see methods).

To establish whether participants can infer the existence of community structure and use a prior over the structural forms to inform their behaviour, we constructed a second experiment. In this experiment, participants were also segregated into two groups. As before, one group learned two hexagonal graphs and the other group learned two graphs with community structure during the first day. However now, both groups learned from random walk and navigate on a community-structured graph during the second day (Figure 1a). Participants who learned graphs with underlying community structure on the first day indeed performed better on the second-day navigation task (number of steps to the target is shorter, Figure 5b upper panel, *D*_*t*=0_, is the initial distance between starting picture and the target, see methods for complete statistical values). Furthermore, they learned the associations faster. While learning the associations participants determined their own learning pace by choosing when to observe the next picture pair. Participants who expected a graph with underlying community structure spend less time on learning each pair of pictures than participants who expected a Hexagonal graph (Figure 4b upper left panel, p=0.003, t=3.19, df = 38, two-tailed ttest). These results suggest that participants’ behaviour was affected by structural prior; the previously experienced graph structure affected the learning policy of participants. The learning policy that is adjusted to underlying community structure leads to faster learning and better task performance. One likely possibility is that, instead of learning the individual pairwise associations, participants simply inferred the community structure and assigned each node to the current community, while identifying the connecting nodes.

In order to understand how different behavioural policies lead to different performance, we examined participants’ choices during navigation. During the navigation part of the task, participants had to choose between two pictures (to get closer to the target) or skip and sample a new pair (if they thought both pictures took them further away). We examined participants’ choices during all trials in which one picture was a connecting node and the other was not. Participants who had the correct prior chose connecting nodes significantly more than participants who had the wrong prior (p=0.015, t = 2.25, df = 38 one-tailed ttest, Figure 5b left lower panel). Furthermore, this cannot be driven by better inference of graph distances as they chose connecting nodes more frequently, *even if this choice was the wrong choice* (p=0.0032, t = 2.88, df = 38, one-tailed ttest, it took them far away from their target, Figure 5b right lower panel). These results imply that participants with the correct structural prior behave according to a policy of ‘prefer connecting node’.

Here, inference of structural knowledge can lead to the transfer of two different types of knowledge. First, the transfer of the abstract transition structure (as participants did in the previous task). Second, the transfer of unique behavioural policy that is tailored to that particular structural form. We cannot identify here whether our results originated from policy transfer only, or whether participants also transfer abstract knowledge of the transition structure itself. According to our model, inference of structural knowledge allows transfer of the relevant basis set. Basis set for community structure enables inference of connecting nodes’ identity immediately when learning the emission matrix. Therefore, exploiting this basis during the task should enable faster identification of these nodes. This implies that transfer of structural knowledge itself (at the form of the relevant basis set) will lead to better identification of these special nodes.

Further, basis set representation generalizes over all tasks that are governed by the same structural form, therefore, they allow the semantic understanding of connecting nodes and the generalization of such policy. This experiment supports this idea but other types of representation for connecting node identity may enable the same behaviour. Being trapped in a community for a long period can be frustrating. Hence, escaping from a community might be perceived as rewarding. One possible explanation for the choice of connecting nodes is therefore simply that they have been more rewarding in the past. However, this cannot explain the effect we observed. We compared the behaviour of two groups of participants that did the exact same task and only differ by the underlying graph structures that have been learned the day before. Participants with the wrong prior spent longer within communities and would, under this argument, experience greater reward when escaping them. Therefore, the value of connecting nodes assigned by the participants with the wrong prior should be higher. If participants chose according to value difference, we should see the opposite effect. This model-free effect therefore runs opposite to the behaviour we observe.

## Discussion

We have shown, using a graph learning task, that participants are able to transfer abstract strcutral knowledge. They were able to transfer abstract transition structure of the task and structurally relevent behavioural policy. They exploited the transfered structural knowledge to infer the existance of unobserved trajectories, identify important task states and improve preformance on the task. Using a computational model we have suggestd a representation for structural forms that allows generalization over particularities of the current task and enables transfer of abstract structural knowledge. Each structural form is represented by a particular basis set that encodes flexibly the transition sturctures that belong to that form. Each basis vector can be streched and compressed according to the infered graph size, hence, the set enables compressed and generalizable representation of the transition structure ^8^. We showed that having such representations enable correct inference of the structural form that govern the associations between states in the task. Approximating the current transition matrix using a basis set allows inference of routes that have not been taken before and inference of the identities of important task states. Our current experimental results suggenst that human do exploit abstract structural knowledge, but whether they achieve this via basis representations requires further experiments.

The ability to transfer structural knowledge requires that participants infer or acquire the correct representation of the hidden structure of the graph during the first day. One possibility is that structural knowledge is acquired slowly via experiencing different scenarios that share the same underlying structural form during our life ^28–30^. In the context of our task, we suggest that the experience during the first day shapes the prior over structural forms on the second day. Then, particpants with the correct structural prior are able to infer the correct structural form faster, estimate better the current transition matrix and transfer the correct behavioral policy, therefore achieveing better preformance in the task.

In the current work we considered two types of structural forms to introduce the idea of basis sets representation for structural knowledge. Theoretically, the idea of basis sets representation for structural knowledge can be extended to other structures that are common in nature, such as rings and hierarchies. For example, eigenvectors of transition matrices with underlying structural form of hierarchy consists the information on the layer of nodes in the hirerchy (see supplementary). Similarly to connecting nodes in a graph with a community structure, this information can be beneficial in preforming a task and allows the learning of the meaning of each layer over a variety of tasks with the same underlying structural form.

The use of basis sets is inspired both by spectral graph theory ^8,15,31^ and by existing research on the hippocampal – entorhinal system. Entorhinal cortex consists of cells that have hexagonal activity pattern while the animal walks freely in an arena (grid cells) ^32^. This activity pattern resembles the patterns of transition matrix eigenvectors of hexagonal graphs ^33,34^. Further, grid-like representations emerge in enviroments/tasks that are not spatial but share similar statistical structure ^35–37^. These observations may suggest that grid cells can be used as basis functions for all environments in which the associations between the states are governed by the rules of 2D Euclidean space.

It has been suggested that grid cells are created using attractor neural networks ^38,39^. This suggestion is supported by the observation that their activity correlation pattern is maintened during sleep ^40,41^. Further, grid cells remap (their activity pattern is shifted) in new environments but their Hexagonal activity pattern remains ^42^. These observations may suggest that grid cells activity pattern is stably represented and therefore can be recalled in new environements that share the same underlying statistical pattern of transition structure, in accurdance with our basis sets hypothesis. Further, our hexagonal graphs were a torus, such that there were no boundaries. Introducing boundaries and a policy in which the agent prefers to stay near the boundaries, similarly to animals behaviour, creates an asymetrical transition structure. The right eigenvectors of such representation hold the variation in transitions and have patterns that reseamble EC bounday cells ^43^ (in addition to hexagonal patterns, see supplamentary). We can think of a boundary cell as part of a basis set that captures special nodes in translational invariants graphs.

Inspired by graph theory, reinforcement learning and the activity patterns in the hippocampal formation, we suggest, that the brain may represent structural froms in a form of basis sets. Using modeling, we show that such basis sets allows transfer of structural knowledge that is relevant to the current task. Our behavioral experiments demonstrate that humans can transfer abstract structural knowledge and exploit it in a new task.

## Methods

### Model description

#### The generative model

In this work, we follow Tenenbaum et al. ^13^ in suggesting that humans represent structural knowledge as structural forms. Each structural form is a family of graphs in which the nodes of the graph are organized according to a particular rule. For example, hexagonal grid connectivity pattern will always present a translational and rotational symmetry, or in community structure, the nodes within a community will be highly interconnected, while the connectivity between communities is sparse. We assume that on each new task, humans infer the structural form that best fit the graph of the current task. Following this inference, they can transfer the relevant information that represents this structural form.

Following Kemp and Tenenbaum ^4^ we formalized the inference over structural forms using hierarchical generative model of graphs. In our task, the observations of the agents and participants are Markovian (each state depends only on the state before) and follow a transition matrix that is characterized by an underlying graph structure. Each graph belongs to one of the structural forms that are considered in our experiments (hexagonal grid and community structure). We assume that each task structural form (*S*_*f*_) is generated by sampling from a uniform distribution over the structural forms. Given a structural form, the graph dimensions (*θ*) are sampled from a prior distribution that is unique for each structural form (see below). Together, *S*_*f*_ and *θ* fully determine the transition matrix of the graph 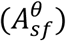. Then, given a transition matrix the emission matrix (*B*) is sampled (in the following, we will not find the posterior of *B*, therefore we do not state any prior for *B* here). Using these two matrices, the observation (*O*) can be generated.

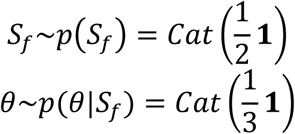

#### Model inversion

The task of the agent is to infer the hidden states of this generative model; given a set of observations, the agent should infer, using Bayes rule, the structural form and graph dimensions (*θ*) that characterized the graph of the current task or environment.

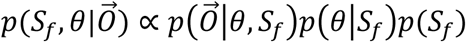

Where:

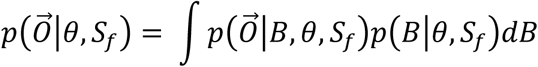

Here the integral is over all possible values of the entries in the emission matrix B. As solving this integral is hard, we have approximated it by using the Bayesian Information Criterion (BIC), therefore:

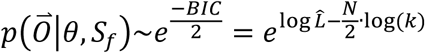

Where *N* is the number of states in the graph, *k* is the number of observations and 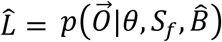 is the likelihood of the sequence of observation with 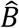 as the Maximum Likelihood Estimate of the emission matrix. As the transition matrix (*A*) is fully defined by the structural form and the dimension of the graph, and the observations depend directly on the transition and emission matrices only, we can write the likelihood as: 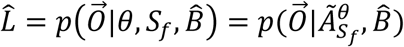 where 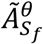 is an estimated transition matrix (see below). Our model is a Hidden Markov Model (HMM), therefore, we can exploit a variant of the Baum-Welch algorithm ^44^ to estimate *B* from the observations and calculate the likelihood 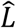 (see supplementary for details). The Baum-Welch algorithm gives a maximum likelihood estimates for the transition and emission matrices as well as the likelihood itself. Here, instead of learning the transition and emission matrices from the data (*O*), we learned only the emission matrix and assumed that the transition matrix is known; for each structural form and graph size that were considered, we approximated the transition matrix using the relevant basis set (see below). For each approximated transition matrix we estimated 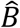 and 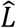.

Using these quantities, we estimated 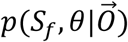 and inferred the current structural form and dimension.

The structural form of the current task is inferred by calculating the posterior and choosing its maximum (MAP):

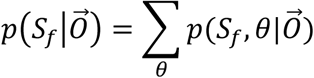

Following the inference of the structural form the current graph size is inferred using MAP of:

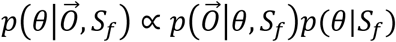

#### Approximating the transition matrices using Basis sets for structural knowledge

To allow generalization over particularities of the current graph structure such as its dimensions (θ), the transition matrices are approximated using basis sets 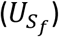 for structural knowledge. Each structural form is represented by unique basis set and transition matrices of all graphs that share structural form are approximated using this set (see below the definitions of the basis set for each of the structural forms). Instead of learning the transition and Emission matrix from the data 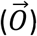, we infer the transition matrix under the assumption that the task transition matrix can be approximated by those basis sets that are already known. Therefore, once the agent solved the inference problem over structural forms and graph size, it can use its prior representations of possible basis sets to estimate the new transition matrix.

For each structural form, given a particular graph size, the basis vectors in the set can be stretched and compressed, using interpolation to adjust for the currently estimated graph. The approximated transition matrix becomes: 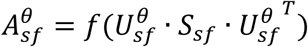, where 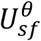 is the adjusted basis set, *S*_*sf*_ is a diagonal matrix of weights (eigenvalues in the hexagonal grid graphs and ones in the community stucture graphs), and *f* is a threshold linear function. We then substract the diagonal and normalized the matrix (see figure 3D).

#### Inferring graph size

We assumed, for simplicity, that there are three different possible graph sizes for each structural form, therefore the prior probabilities of θ are uniform within this set and zero otherwise. We further assumed, for simplicity, that the two dimensions of the hexagonal grid are equal and the number of nodes in a community is also equal, hence θ defines a vector of possible number of nodes in a graph. For hexagonal graphs we considred N = [25,36,49], for a graph with underlying community structure N = [28,35,42] with equal prior probability. We emphasis that the basis set 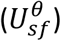 for each transition structure within a structural form is a scaled or trancated version of a general basis set for that structural form.

#### Estimating distances between two pictures

Using the inferred graph transition and emission matrices, the agent approximated the distance between two observations. As the transition structure is approximately known, we estimated the distance matrix between two abstract states on the graph using this approximation; we adopted a threshold function of the transition matrix to estimate the Adjacency matrix and then estimated the abstract distance matrix (*D*(*z*_*m*_, *z*_*k*_)) using it. As the distance matrix represents the distances between abstract states (*z*_*l*_), the distances or the number of steps between the observations themselves is calculated by:

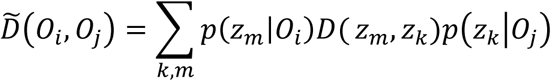

The emission matrix gives us 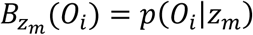, as *p*(*z*_*m*_) is uniform, we inverted *p*(*O*_*i*_|*z*_*m*_) by:

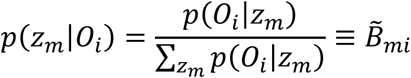

Then, we multiply the abstract distance matrix by the 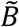 to get the distance matrix between the observations:

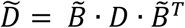

Using this matrix, we have calculated the agent answers in choosing between pictures (observations), the picture that is closer to the target picture. We would like to note here that the actual emission matrix in our task is not probabilistic and it is an identity matrix. When the agent estimates the emission matrix from the observations it converges to any permutation matrix that keeps the symmetry of the graph. There are other approximations for the distances between observations that can be adopted which take into account the probability structure of the transition matrix, such as the Successor Representation ^24^.

### Connecting node inferece

For estimating the number of stimuli that are correctly inferred as connecting nodes we calculated for each simulated block the fraction of correctly inferred connecting node as: 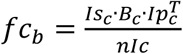 where *Is*_*c*_ is a vector at the size of the inferrd number of states, with one for a connecting node state and zero otherwize, *Ip*_*c*_is a vector at the size of the number of stimuli, with one for a stimuli of connecting nodes and zero otherwize. *nIc* is the number of connecting nodes in the inferrd graph size. The fraction of incorrect inferred connecting node is defines as: 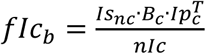 where *Is*_*nc*_ = 1 − *Is*_*c*_.

#### Basis sets

### Community structure

The number of nodes within each community is considered constant for simplicity. There is an assignments vector for each community with one for each node that belongs to that community and zero otherwise. The number of such vectors is determined by the number of communities that are currently considered. Further, there are ‘connecting node assignemnt vectors’ for each community, which gives a probability for a node in a community to connect to another node in another community, the probability for a second connecting node is lower.

### Hex

The eigenvectors of large hexagonal graph were computed. We kept as a basis sets the 12 most informative eigenvectors (excluding the constant). We then resize the eigenvectors according to the the size of the graph that is currently considered using standard interpolating method.

### Experimental details

#### Behavioral experiments

Experiment 1: transfer of Hexagonal structure: 30 participants in each group.

Experiment 2: transfer of community structure: 20 participants in each group.

Participnats learned two graphs with the same underlying structure but different stimuli during the first day. Stimuli were selected randomly, for each participant, from a Bank of stimuli (separate Bank for each graph). Each graph was learnt in four blocks (figure 1b). A third graph was learnt on the second day during seven blocks of the task. During the first day, in the two experments, participants learned the graph by observing pairs of stimuli that were taking from random walk on the graph. When we examined knowledge transfer of community structure (second experiment), participants learned the graph using random walk during the second day as well.

##### Examining transfer of Hexagonal structure

A link that is constantly missing may lead to an inference of the existence of an obstacle. If participants infer the existence of an obstacle, they will not infer the existence of an unobserved link. We speculated that learning by sampling pairs of neighbouring nodes, instead of learning from pairs that are taken from random walk on the graph, would reduce the inference of borders and obstacles. Reducing the inference of borders and obstacles is an important requirement for our experiment; otherwise, participant will not infer the existence of a link. Following the same reasoning, we have excluded the navigation task during the second day of that experiment.

#### Experimental details

4 blocks for each map on the first day. 7 blocks on the second day (third map).

##### Learning phase

Random walk: Hexagonal graphs: 120 pictures per block. Community structured graph: 180 pictures per block (more pictures as in a random walk the smapling per community is very large). Pairs: 150 pairs in each block.

##### Maximum steps in the navigation task

200. The trial has stopped when a participant exceed this number and a massage that calims ‘to many steps’ has been shown. *Number of questions in the pile task and ‘is it in the middle’* task were 16 in each block.

##### In the estimating distance questions

45 questions per block.

#### Statistical values – second experiment

Correct strutural prior leads to faster navigation to the target:

Number of steps to the target is two, p-value = 0.00095, t = 3.33

Number of steps to the target is three, p-value = 0.025, t = 2.02

Number of steps to the target is four, p-value = 0.0077, t = 2.54 One-tailed ttest, df = 38

## Acknowledgements

We thanks Alon Baram for careful reading of the manuscript and Tamas Madaresz for his support in addressing theoretical questions.

## Author contributions

S.M. and T.B. conceived the study. S.M. and T.B. designed the experiment. S.M. programmed the experiments. S.M. performed the experiments. S.M. and T.B. developed the models. R.M and T.P advise on the model. S.M implemented the model. S.M. and T.B wrote the manuscript. All authors discussed the results and commented on the paper.

