## Supplementary for "Transferring structural knowledge across cognitive maps in humans and models"

**Estimating $p(O|A_{sf}^{\theta}, \hat{B})$ and B using a variant of the Baum-Welch algorithm:**

*Learning from Randm walk*

For each considered graph dimention, that belongs to one of the structural forms, we have estimated the transition matrix ($A_{sf}^{\theta}$) using the rescaled basis set of that from. We then estimated, for each approximated transition matrix, the emission matrix B and the probability of the observations given these matrices, $p(O|A_{sf}^{\theta}, \hat{B})$, using the Baum-Welch algorithm. The Baum-Welch algorithm estimates both the emission matrix and the transition matrix, here, we assumed that the transition matrix is known and therefore did not estimate it. We addapted the matlab routin 'hmmtrain' such that the transition matrix is given and not learned and estimated B and $p(O|A_{sf}^{\theta}, \hat{B})$.

Simulations details: During our simulations the agents saw 150 pairs of pictures within each block.

When the correct underlying structure was of a community structure we averaged our results over 20 simulations, while when it was Hexagonal grid we averaged over 15 simulations.

Intialization of parameters:

*Hexagonal grid*:

$$B_{z=1}(o=1) = 1,\ B_{z=1}(o \neq 1) = 0, B_{z\neq 1}(o=1) = 0, B_{z=i}(o=j) = p_0\ i,j \neq 1$$

$$p_0 = \frac{1}{N_p - 1}$$

Where *z* is a state and *o* is an observation, $N_p$ is the number of observations.

*Community structure*:

Because of the symetry of the community graph structure, the first picture (observation) can be a connecting node (therefore will assign arbitrarily to state 1) or a 'not connecting node' and therefore assign arbitrarily to state 2. As there are 2 connecting nodes in a community the initial condition to the emission matrix is:

$$B_{z=1}(o=1) = \frac{2}{N_c}, B_{z=2}(o=1) = \frac{N_c - 2}{N_c}$$

$$B_{z=1}(o \neq 1) = p_0 \frac{N_c - 2}{N_c}, B_{z\neq 1}(o=1) = \frac{2p_0}{N_c}, B_{z=i}(o=j) = p_0\ i,j \neq 1$$

Where $N_c$ is the community size.

The simulations were intiated in state 1.

*learning from pairs (EM)*

Learning from pairs that are randomly sampled from the graph, istead of a random walk (in which each pair has a picture from the previous pair), results in a process which is HMM of length 2. Therefore, using the Baum-Welch is not relevant anymore. Instead, we estimated the emission matrix ($B$) and the likelihood, $p(\vec{O}|A_{sf}^{\theta},\hat{B})$, by the following EM algorithm:

Given a structural form and a graph dimension the transition matrix is approximated as before ($A_{sf}^{\theta}$). For each particular transition matrix we calculated the following EM steps until convergence:

(1) Expectation step:
In the following, $\vec{z}$ denotes the hidden states. $B^{s-1}$ is the estimation of the emission matrix from the previous EM step (where $s$ is the step).

The observation of each pair of pictures is independent from the previous pair, therefore the probability of the observations and hidden states is given by:

$$p(\vec{O},\vec{z}|A_{sf}^{\theta},B)=\prod_{n} p(O_1^n,O_2^n,z_1^n,z_2^n|A_{sf}^{\theta},B)$$

Where $o_1^n$ ($o_2^n$) is the first (second) observation in the *n's* pair, and $z_1^n$ ($z_2^n$) are the states of the first and second observation respecilvly.

Lets define:

$$\begin{aligned}\alpha_n(z_1^n=i,z_2^n=j)&\equiv p(O_1^n,O_2^n,z_1^n=i,z_2^n|A_{sf}^{\theta},B)\\&=p(z_1^n=i)\cdot p(O_1^n|z_1^n=i,B)\\&\cdot p(z_2^n=j|z_1^n=i,A_{sf}^{\theta})p(O_2^n|z_2^n=j,B)=\frac{1}{N}B_{z_1^n}(o_1^n)A_{ij}B_{z_2^n}(o_2^n)\end{aligned} \quad (1)$$

$N$ is the number of nodes on the graph. Each pair is sampled randomly and unifrmly therefore: $p(z_1^n)=\frac{1}{N}$. $B_{z_k^n}(o_k^n)=p(O_k^n|z_k^n)$ and $A_{ij}=p(z_2^n=i|z_1^n=j)=\begin{cases}\frac{1}{6} & if\ i\&j\ are\ neighbours\\ 0 & otherwise\end{cases}$.

and:

$$\begin{aligned}\log(p(\vec{O},\vec{z}|A_{sf}^{\theta},B)&=\sum_n \log[p(O_1^n,O_2^n,z_1^n,z_2^n|A_{sf}^{\theta},B)]=\sum_n \log[\alpha_n(z_1^n,z_2^n)]\\&=\sum_n\left(-logN+logB_{z_1^n}(o_1^n)+logA_{ij}+logB_{z_2^n}(o_2^n)\right)\end{aligned}$$

The expected log liklihood of the data $(\vec{O},\vec{z})$ given the current model parametrs $(A^{\theta}_{sf})$, the previouse estimation of the parameters $(B^{s-1})$ and the the distribution of the states $(\vec{z})$ conditioned on observation $(\vec{O})$ becomes:

$$Q(B,B^{s-1}) = \sum_{z\in Z} p\big(\vec{z}|\vec{O},A^{\theta}_{sf},B^{s-1}\big)\log(\,p\big(\vec{O},\vec{z}\big|A^{\theta}_{sf},B\big)$$
$$= \sum_{z\in Z} p\big(\vec{z}|\vec{O},A^{\theta}_{sf},B^{s-1}\big)\sum_{n}\log[\alpha_n(z_1^n,z_2^n)]$$

And the liklihood of the current estimation of *B* is:

$$L(B^{s-1}) = \; p(\vec{O}|A^{\theta}_{sf},B^{s-1}) = \prod_{n} p(O_1^n,O_2^n|A^{\theta}_{sf},B^{s-1})$$
$$= \prod_{n}\sum_{i,j}\alpha_n^{s-1}(z_1^n=i,z_2^n=j)$$

(2)

Where $\alpha_n^{s-1}(z_1^n=i,z_2^n=j) = p\big(O_1^n,O_2^n,z_1^n=i,z_2^n=j|A^{\theta}_{sf},B^{s-1}\big)$

On each expectation step we calculated: $\alpha_n^{s-1}(z_1^n=i,z_2^n=j)$ and $L(B^{s-1})$.

(2) maximization step:

We define a Lagrangian ,using Langrange multiplier to satisfy the condition that $\sum_k B_i(k)$=1:

$$\hat{L}_{\lambda} = Q(B,B^{s-1}) - \sum\nolimits_i \lambda_i\left(\sum\nolimits_k B_i(k)-1\right)$$

As we are maximizing in respect to *B*, we can keep only the the following terms:

$$\tilde{L}_{\lambda} = \sum_{z\in Z} p\big(\vec{z}|\vec{O},A^{\theta}_{sf},B^{s-1}\big)\sum_{n}\left(logB_{z_1^n}(o_1^n)+logB_{z_2^n}(o_2^n)\right) - \sum\nolimits_i \lambda_i\left(\sum\nolimits_k B_i(k)-1\right)$$
$$= \sum_{i,j}\sum_{n} p\big(z_1^n=i|\vec{O},A^{\theta}_{sf},B^{s-1}\big)\,logB_i(O_1^n) + p(z_2^n$$
$$= j|\vec{O},A^{\theta}_{sf},B^{s-1})logB_j(O_2^n) - \sum\nolimits_i \lambda_i\left(\sum\nolimits_k B_i(k)-1\right)$$

Maximizing in respect to $B_i\big(O_{1,2}^n=k\big)$ gives:

$$\frac{\partial\hat{L}_{\lambda}}{\partial B_i(k)} = \frac{\sum_n\big(p\big(z_1^n=i|\vec{O},A^{\theta}_{sf},B^{s-1}\big)\delta(o_1^n=k)+p\big(z_2^n=i|\vec{O},A^{\theta}_{sf},B^{s-1}\big)\big)\delta(o_2^n=k)\big)}{B_i(k)}$$
$$-\lambda_k$$

For solving for $B_i(k)$ we define:

$$\tilde{\gamma}_n(i,j) \equiv p\left(z_1^n = i, z_2^n = j \middle| \vec{O}, A_{sf}^{\theta}, B^{s-1}\right) = \frac{p(z_1^n(i), z_2^n(j), \vec{O} | A_{sf}^{\theta}, B^{s-1})}{p(\vec{O} | A_{sf}^{\theta}, B^{s-1})}$$

$$= \frac{p(z_1^n = i, z_2^n = j, O_1^n, O_2^n | A_{sf}^{\theta}, B^{s-1})\, p(O_{\sim n} | A_{sf}^{\theta}, B^{s-1})}{p(\vec{O} | A_{sf}^{\theta}, B^{s-1})}$$

$$= \frac{\alpha_n^{s-1}(z_1^n = i, z_2^n = j)\, p(O_{\sim n} | A_{sf}^{\theta}, B^{s-1})}{p(\vec{O} | A_{sf}^{\theta}, B^{s-1})}$$

Where:

$$p(O_{\sim n} | A_{sf}^{\theta}, \hat{B}) = \prod_{m \sim n} \sum_{i,j} \alpha_m(z_1^m = i, z_2^m = j)$$

Therefore:

$$\gamma_n(i) \equiv p\left(z_1^n = i | \vec{O}, A_{sf}^{\theta}, B^{s-1}\right) = \sum_j \tilde{\gamma}_n(i,j)$$

Note that $\alpha_n^{s-1}(z_1^n = i, z_2^n = j)$ and $p(\vec{O} | A_{sf}^{\theta}, B^{s-1})$ were calculated in the Expextation step.

Solving for $\frac{\partial \hat{L}_\lambda}{\partial B_i(k)} = 0$, plug in $\gamma_n(i) \equiv p\left(z_1^n = i | \vec{O}, A_{sf}^{\theta}, B^{s-1}\right)$ and normelizing gives:

$$B_i^s(k) = \frac{\sum_{n=1}^{T} [\delta(o_1^n = k)\, \gamma_n^1(i) + \delta(o_2^n = k) \gamma_n^2(i)]}{\sum_{n=1}^{T} [\gamma_n^1(i) + \gamma_n^2(i)]} \tag{3}$$

Therefore on each step we calculate $\gamma_n(i)$ given $\alpha_n^{s-1}$, and use equation (3) for the new estimation of the emission matrix ($B^s$).

These 2 steps repeat until convergence. We then estimate $\log L(B)$ using the estiamted $B$.

Here, during our simulations the agents saw 150 pairs of picture within each block.

*Intiation of B:*

*Hexagonal grid*:

$$B_{z=1}(o = 1) = 1, B_{z=1}(o \neq 1) = 0, B_{z \neq 1}(o = 1) = 0,$$

$$B_{z=2}(o=2)=1, \qquad B_{z=2}(o\neq 2)=0, B_{z\neq 2}(o=2)=0, B_{z=i}(o=j)=p_0\ i,j\neq 1,2$$

$$p_0=\frac{1}{N_p-2}$$

*Community structure*: stayed as above.

The results here are averaged over 30 simulations.

***Supplementary figures:***

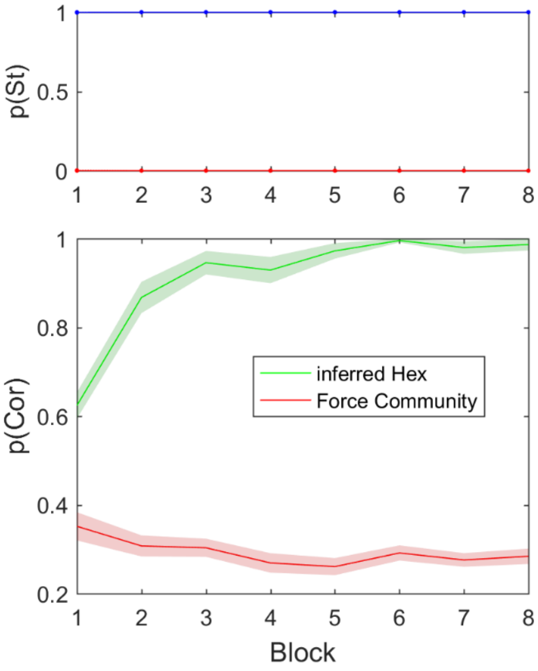


***S.Fig1 Learning from pair - simulations***

*Upper panel - When learning from pairs with missing link, the model could infer the structural form correctly. Blue p(Hexagonal grid), red p(community structure).*

*Lower panel - while inference of the correct structural form leads to high above chance performance in detecting which of two pictures is close to a target picture (green), having only the wrong structural form information (only community structure basis set are available), leads to performance which is significantly less than chance (red).*

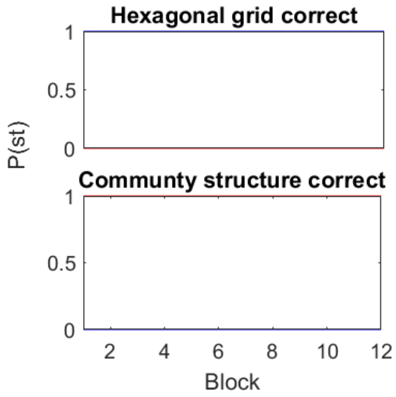


**S. Fig2: Inference of correct structural form was immediate.**

Blue: probability of Hexagonal structural form. Red: probability for community structural form. Agent learned from random walk.

Hex correct: Cluster correct:

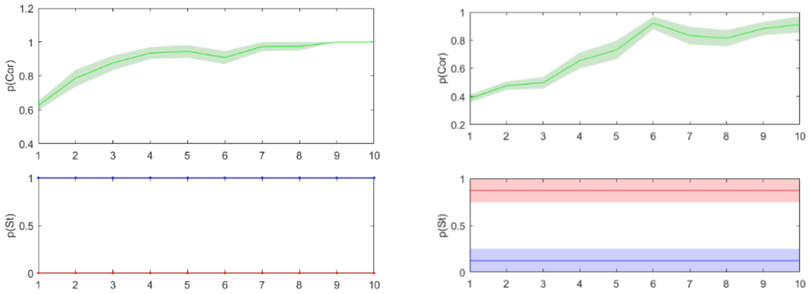


**S.Fig3: Inference on larger and equal size graphs**

**49x49 graphs**: as the graph sizes were not completely identical we simulated 49 nodes Hexagonal and community structure graphs to show that structural inference was not biased by graph size. Here the agents learned from random walk.

Border cells:

Entorhinal cortex, the area in which grid cells exist, also has border cells. These cells are active when the animal is near a border. Here we show that changing the behavioural policy of the agent from random walk/ uniform over the graph, into a policy that ‘prefers’ staying near the borders results in transition matrix eigenvectors that both resemble grid (S. figure 4b) cells but also border cells (S. figure 4a). This suggests that border cells, similarly to grid cells, may be part of a basis sets to represent tasks on translational invariants graphs (with borders).

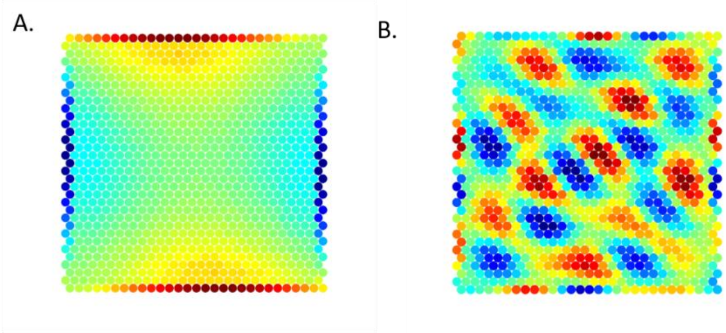


**S. Fig4: Eigenvectors of a transition structure with a Policy of ‘prefer borders’ have EC border cells pattern.**

Here we simulate the wish to stay near a boundary by increasing the probability to move to a node on the boundary to be 15 times larger than to move out of the boundary.

A. An example for an eigenvector with boundary cell pattern
B. An example for an eigenvector with hexagonal grid pattern.

Basis set for representing hierarchical structure**:**

In the main manuscript, we discussed mainly two structural forms; hexagonal grid and community structure. The idea of basis sets can be extended to any other structural form. Here we show that eigenvectors of hierarchical graph transition matrix have interesting patterns that encode the layer information on the hierarchical graph. Similar to abstract representation of connecting nodes and borders, abstract representation of the hierarchical graph layers can support the understanding of the semantic meaning of the layers and the node at the head of the hierarchy.

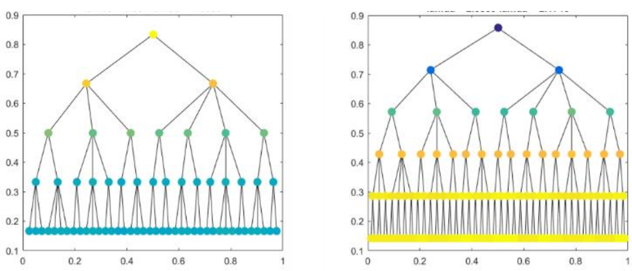


**S. Fig5: Hierarchical graph transition matrix eigenvectors that represent layers information:**

Here we show two examples of eigenvector that represent the distance from the head node.

Human results in learning associations on the graph

Participants knowledge of the associations were much better than chance (tasks part 2 & part 3 p<<0.001) although the graphs were very large (35, 36 nodes) with degree of 6.

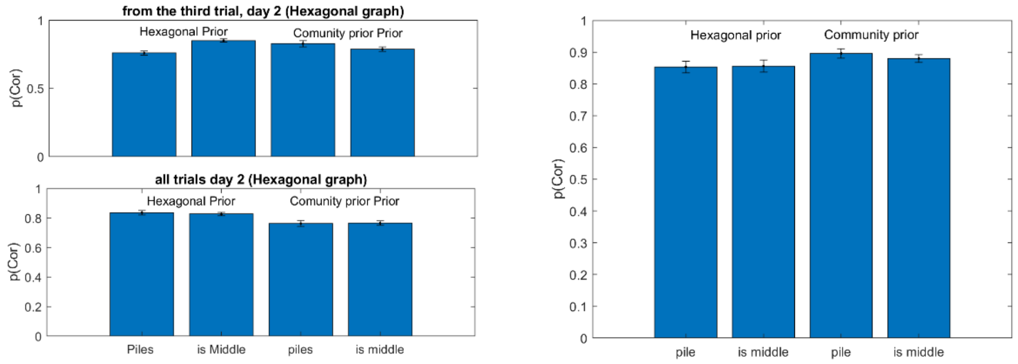


**S.Fig6: Learning pairwise associations: (questions type 2 and 3 in each block), second day**

A) pairwise associations knowledge, first experiment. When including the first 2 blocks (out of seven) the participants with Hexagonal prior perform better on the associations task, this different disappear after the second block.

B) pairwise associations knowledge, second experiment
